# Pathological crosstalk between AQP4-dependent ATP/Adenosine release and dopamine neurotransmission underlying depressive behavior during cocaine withdrawal

**DOI:** 10.1101/2022.10.03.510559

**Authors:** S. Okada, M. Kobayashi, H. Lee, C. Nuralifah, M. Morita

## Abstract

Our previous study has demonstrated the AQP4-dependent ATP release and following elevation of extracellular adenosine in hippocampal slices during hypo-osmotic treatment. In the present study, we examined the crosstalk between AQP4-dependent adenosine elevation and dopaminergic neurotransmissions, especially focusing on the mouse depressive behavior during cocaine withdrawal. The depressive behavior was mitigated in AQP4 knockout mice, and was sensitive to adenosine A_1_ receptor antagonists. Thus, the influence of cocaine withdrawal on the adenosine or dopamine elevation in the slices from the striatum or medial prefrontal cortex, which is the focus of depression was analyzed by the originally-developed biosensor cells, which elevates intracellular Ca^2+^ in response to adenosine or dopamine. We have found that the evoked-dopamine release in the striatum is tonically suppressed via A_1_ receptor in a AQP4 dependent manner, and this suppression is augmented by the cocaine withdrawal. In contrast, the evoked dopamine elevation in the medial prefrontal cortex was suppressed by treating slices with adenosine or an A_1_ receptor antagonist, and the cocaine withdrawal abolished the suppression by A_1_ receptor antagonist alone in an AQP4-dependent manner. Since a GABA_A_ or group II metabotropic glutamate receptor antagonist occluded the suppression by A_1_ receptor antagonist, it has been suggested that the glutamatergic and GABAergic terminals suppressing dopamine releases have A_1_ receptors, which the cocaine withdrawal down-regulates in a AQP4-dependent manner. These results indicate that astrocytes modulate dopaminergic neurotransmissions via A_1_ receptor in an AQP4-dependent manner, and suggest that cocaine causes depression by changing this modulation.

## Introduction

Adenosine is a neuromodulator, which mediates presynaptic inhibition by A_1_ receptor and synaptic plasticity by A_2A_ receptor (Chen et al., 2001). Despite the associations of extracellular adenosine concentration in the brain with circadian cycle (*) and pathological conditions including stroke (*) and epilepsy (*), the pathways for adenosine release in the brain are not fully determined. Our previous study using a biosensor cell line, which increases intracellular Ca^2+^ in response to adenosine for measuring the adenosine release in hippocampal slices has demonstrated two novel distinct pathways, namely the calcium channel-dependent neuronal adenosine release following electrical stimulation and the AQP4-dependent ATP release and subsequent adenosine elevation due to ATP degradation following hypoosmotic treatment (*). AQP4 is a water channel exclusively expressed in astrocytes in the brain (*), thus hypoosmotic treatment likely induces the astrocytic ATP release due to water influx and subsequent regulatory volume decrease (*). This pathway was further supported by our related study showing the involvement of volume-regulated anion channel in the ATP release from cultured astrocytes following mechanical stimulation (*).

Accumulating evidence indicates antagonistic interactions between adenosine and dopamine in the brain. A_2A_ and D_2_ receptors form heterodimer and suppress each other (Hillion et al., 2002), while A_1_ receptor inhibit dopamine release in the striatum and spinal cord (Acton et al., 2018; Ross and Venton, 2015). The upregulation of dopamine neurotransmission by adenosine receptor antagonist is a clinical strategy for treating Parkinson’s disease patients (Kondo and Mizuno, 2015). In addition, chemicals, which act on adenosine receptors, as well as modulate extracellular concentration of adenosine have been shown to affect schizophrenia (*) or bipolar disorders (*), which involve pathological changes of dopaminergic neurotransmission. Furthermore, astrocytes likely participate in the crosstalk between adenosine and dopamine, because it has been reported that the A_1_ receptor activation by the adenosine from astrocytes tonically inhibit dopamine release (https://pubmed.ncbi.nlm.nih.gov/35042768/), as well as is associated with the anti-depression effect of sleep deprivation (https://www.ncbi.nlm.nih.gov/pmc/articles/PMC3566717/). Thus, the astrocyte atrophy in depression patients (Zhao et al., 2018) may cause symptoms by changing extracellular adenosine, as in the case of epilepsy, which is developed as a consequence of reduced extracellular adenosine due to the pathological change of astrocyte (Fedele et al., 2005).

The present study has hypothesized that the changes of crosstalk between the AQP4-dependent adenosine elevation and dopaminergic neurotransmissions underlie the mouse depression-like behavior during cocaine withdrawal (Filip et al., 2006). Our approach includes the behavioral analysis of wild type and AQP4^-/-^ mice and the measurement of adenosine or dopamine elevation in brain slices from the foci of depression, the striatum and medial frontal cortex (mPFC) by biosensors as in our previous study (*). Once the association between the AQP4-dependnet adenosine elevation and the pathological changes of dopamine neurotransmission in depression is established, astrocytes and water dynamics in the brain will be recognized as a new diagnostic and therapeutic target of psychiatric disorders.

## Materials & Methods

### Animal experiments

All animal experiments were approved by the institutional animal care and use committee of Kobe University (Permission number: 25-09-04) and performed in compliance with the Kobe University Animal Experimentation Regulations. C57BLC background mice were maintained at 24℃ and under 12h light/dark cycle. Water and food were provided ad libitum. Mice at least 8 weeks old were used for experiment. AQP4 knock out mice (AQP4^-/-^) (Kitaura et al., 2009) were provided by Dr. Mika Terumitsu (National Center of Neurology and Psychiatry, Tokyo, Japan). Depression-like behavior during cocaine withdrawal was accessed by the forced swim test (FST) 14 to 17 days after five consecutive days of intraperitoneal cocaine injection at 15 mg/kg. For the FST, mice were acclimatized in an experiment room with white noise for one hour, then placed in a transparent cylinder 22 cm in diameter filled with water at 24°C at a height of 12.5 cm (Can et al., 2012). The mobility of mice was recorded as 8 min RGB videos at 15 fps by a web camera, C270n (Logicool, Tokyo, Japan) and quantified as described by Gao et al (Gao et al. 2014). Each frame in red color channel was converted to 8 bit gray-scale and subtracted by previous one. Then, the number of pixels with a value above five were counted and averaged for 15 consecutive frames as the mobility of mice in one second, The immobility time was measured as the duration, in which the mobility was below an arbitrary threshold in 60 secconds.

### Biosensor cells

As a biosensor for adenosine or dopamine, a stable CHO cell lines expressing RCaMP, G_qi_5 and mouse A_1_ receptor (A_1_-CHO) or human D2 receptor (D_2_-CHO) were established. CHO cells were maintained in Ham’s F12 medium containing 10% fetal bovine serum and antibiotics (50 U/ml penicillin and 50 µg/ml streptomycin). For [Ca^2+^]_i_ imaging, cells were seeded on coverslips. Clonal cells were obtained by transfecting with pCAG-cyto-RCaMP (Addgene), pME- G_qi_5 (Yamashiro et al***) and pCMV-mA1 (OriGene) or DRD2-Tango (Addgene) by Lipofectoamine3000 and screening with 200 ug/mL G418. Then, the A_1_-CHO or D_2_-CHO was selected as a cell line responding to 100 µM adenosine or 100 µM dopamine, respectively by Ca^2+^ imaging.

### Brain slice preparation

Mice were anesthetized mice by intramuscular injection of ketamine (100mg/kg, Daiiichi-sankyo, Tokyo, Japan) and xylazine (10 mg/kg, Bayer Yakuhin, Osaka, Japan) and decapitated. Brains were extracted, put into an ice-cold cutting solution containing (in mM): 2 KCl, 1.5 NaH_2_PO_4_, 26 NaHCO_3_, 6 MgSO_4_, 0.2 CaCl_2_, 10 D-glucose and 220 sucrose, and saturated with 95% O_2_ and 5% CO_2_. Coronal slices of 400µm thickness were cut using a microslicer (ZERO-1, Dosaka EM, Kyoto, Japan) and maintained in artificial cerebrospinal fluid (aCSF) containing (in mM): 124 NaCl, 2 KCl, 1.5 NaH_2_PO_4_, 26 NaHCO_3_, 1 MgSO_4_, 2 CaCl_2_, 10 D-グルコース, for more than one hour before experiment.

### Calcium imaging

Slices were placed on the biosensor cells and perfused with aCSF at 5 mL/min. The RCaMP images of the biosensor cells were obtained every one second by using an inverted microscope (IX-70, Olympus, Tokyo, Japan) equipped with an objective (UApo/340 20x / 0.70, Olympus) and a cooled-CCD camera (Orca-R2, Hamamatsu Photonics, Hamamtsu, Japan), and F/F_0_ was calculated by ImageJ. Electrical stimulation was given by house-made concentric electrode, isolator (ISO-Flex, Funakoshi, Tokyo, Japan) and pulse generator (Electronic Stimulator, Nihon Koden, Tokyo, Japan). For hypoosmotic treatment, the hypoosmotic aCSF of the osmolarity indicated were prepared by reducing NaCl concentration, and slices were pretreated with the isosmotic low NaCl aCSF, which was supplemented with sucrose in the place of NaCl to achieve iso-osmolarity.

### Reagents

If not specified, reagents purchased from Nacalai tesque (Kyoto, Japan).

### Statistical analysis

For statistical analysis by Excel 2016, 50 biosensor cells were randomly selected from each slice in Fig2BC and 5AC while eight responding biosensor cells were randomly selected in Fig3, 4, 5BD, 6, 7 and 8.

## Results

### The involvement of AQP4-dependent adenosine elevation in the mouse depression-like behavior during cocaine withdrawal

The association of the AQP4-dependent adenosine elevation with the mouse depression-like behavior during cocaine withdrawal was examined by the FST. The immobility times of wild type mice treated with saline (WT / (-)) or cocaine (WT / Cocaine), were respectively 21.33±13.00 sec or 35.82±12.08 sec between 2 and 3 min after starting test, and 19.53±14.09 sec or 36.74±8.59 sec between 3 and 4 min, indicating that the cocaine withdrawal significantly increased the immobility time, thus induced depression-like behavior (Fig. 1A). This depression-like behavior was abolished in AQP4^-/-^ (Fig. 1B). The immobility times of saline (AQP4^-/-^ / (-)) or cocaine (AQP4^-/-^ / Cocaine) treated AQP4^-/-^ mice were respectively 34.77±10.67 sec or 14.83±16.25 sec between 4 and 5 min after stating test, 36.07±10.53 sec or 20.92±15.94 sec between 5 and 6 min, 38.28± 12.06 sec or 20.57±14.10 sec between 6 and 7 min, indicating that the cocaine withdrawal significantly reduced the immobility time, thus failed to induce the depression-like behavior in AQP4^-/-^ mice. Then, the involvement of adenosine in the depression-like behavior was examined by a non-selective adenosine receptor antagonist, caffeine. Saline or 50 mg/kg caffeine was administered to wild type mice during the cocaine withdrawal 40 min prior to the FST. The immobility time of wild type mice during cocaine withdrawal treated with saline (WT / Cocaine / (-)) or caffeine (WT / Cocaine / Caffeine) were respectively 15.64 ± 7.78 sec or 5.74 ± 7.19 sec between 1 and 2 min after starting test, 27.61 ± 6.42 sec or 9.70 ± 7.31 sec between 2 and 3 min, 23.42±5.39 sec or 6.33 ± 8.71 sec between 3 and 4 min, 31.56 ± 9.35 sec or 9.85 ± 11.43 sec between 4 and 5 min, 39.56 ± 8.47 sec or 13.50±10.11 sec between 5 and 6 min, 39.82 ± 10.29 sec or 21.58 ± 13.59 sec between 6 and 7 min, and 44.94 ± 3.37 sec or 19.66 ± 13.22 sec between 7 and 8 min, indicating that caffeine significantly reduced the immobility time as in AQP4^-/-^ mice, thus inhibited the depression-like behavior (Fig. 1C). The effect of caffeine was likely mediated by adenosine A_1_ receptor, because an A_1_-selective antagonist, 8-Cyclopentyl-1,3-dipropylxanthine (DPCPX), which was administered at 4mg/kg, 30 min prior to the FST also abolished the depression-like behavior (Fig. 1D). The immobility time of wild type mice during cocaine withdrawal treated with saline (WT / Cocaine / (-)) or DPCPX (WT / Cocaine / DPCPX) were respectively 6.48 ± 5.94 sec or 0 ± 0 sec between 1 and 2 min, 21.65 ± 11.55 sec or 6.17 ± 7.01 sec between 3 and 4 min, 37.71 ± 5.49 sec or 5.67 ± 6.39 sec between 5 and 6 min, 36.75 ± 14.42 sec or 5.68 ± 7.04 sec between 6 and 7 min, and 40.28 ± 14.14 sec or 20.97 ± 10.60 sec between 7 and 8 min, indicating that DPCPX also significantly reduced the immobility time, thus inhibited the depression-like behavior. These results indicates the involvement of AQP4 and adenosine A_1_ receptor in the depression-like behavior during cocaine withdrawal and suggest that cocaine causes depression by changing the AQP4-dependent adenosine elevation.

**Fig. 1.**
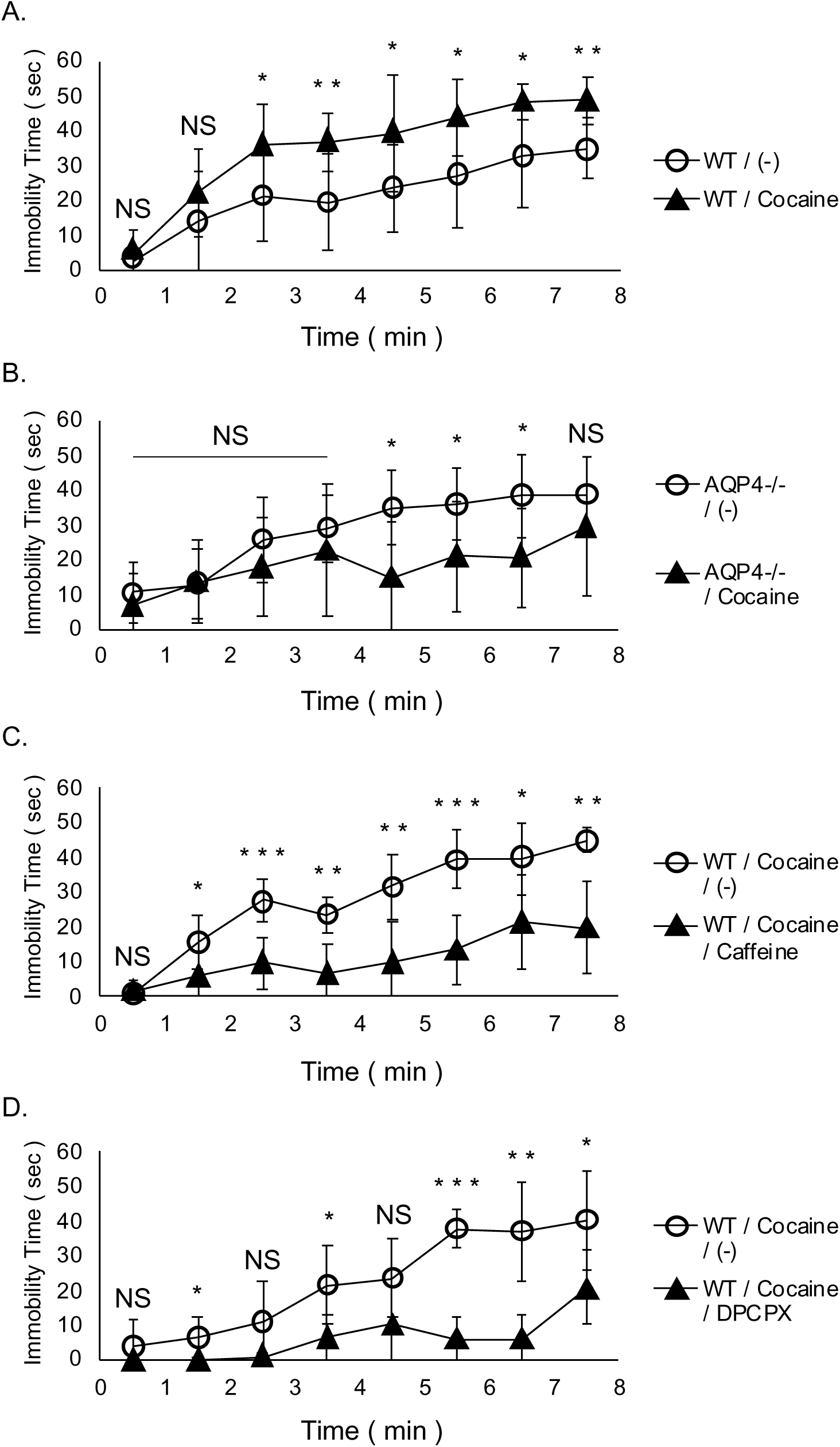
Depressive behavior of cocaine-treated mice were suppressed by adenosine receptor antagonists or in AQP4^-/-^ mice. Depressive behavior during cocaine withdrawal was assessed by the time-dependent changes of immobility time in forced swim test and the influences of pharmacological or genetical interventions were examined. A. Cocaine induced depressive behavior. Wild type mice were treated with saline (WT / (-), n = 8) or cocaine (WT/Cocaine, n=8). Saline or cocaine were treated as described in Materials and Method for inducing depressive behavior during cocaine withdrawal. B. Depressive behaviors of AQP4^-/-^ mice. AQP4^-/-^ mice were treated with saline (AQP4^-/-^ / (-), n = 6) or cocaine (AQP4^-/-^ / cocaine, n = 7). C. The effect of a non-selective adenosine receptor antagonist, caffeine. Saline (WT / Cocaine / (-), n = 6) or 50 mg/kg caffeine (WT / Cocaine / Caffeine, n = 7) were treated 40 min prior to forced swim test. D. The effect of an A_1_ adenosine receptor selective antagonist DPCPX. Saline (WT / Cocaine / (-), n = 5) or 4 mg/kg DPCPX (WT / Cocaine / DPCPX, n = 5) were treated 30 min prior to forced swim test. Mean ± SD. * p<0.05, ** p<0.01, *** p<0.001, NS, not significant.

### The upregulation of AQP4-dependnet adenosine elevation in the striatum during cocaine withdrawal

The adenosine elevation in the ventral part of striatum including nucleus accumbens following electrical stimulation or hypoosmotic treatment was examined by the A_1_-CHO. An electrical stimulation (30Hz, 150 pulses), which had induced adenosine elevations in hippocampal slices (Yamashiro et al., 2017) did not elevate adenosine in striatum slices (Fig2A). Thus, it was suggested that neuronal adenosine release is limited the striatum. Meanwhile, adenosine elevations were detected during a hypoosmotic treatment at -124 mOsm. In addition, the striatal slices from mice during cocaine withdrawal showed 223.14±28.60% significantly larger hypotonically-induced adenosine elevation quantified as the area under the curve (AUG) of Ca^2+^ elevations in the A_1_-CHO, than saline treated mice (Fig. 2B). Furthermore, the upregulation of hypoosmotically-induced adenosine elevation during cocaine withdrawal was occluded in AQP4^-/-^ mice (Fig. 2C). Thus, both the increase of adenosine elevation and the depression-like behavior during cocaine withdrawal were AQP4-dependent, suggesting that cocaine develops depression by increasing the AQP4-dependent adenosine elecvation.

**Fig. 2.**
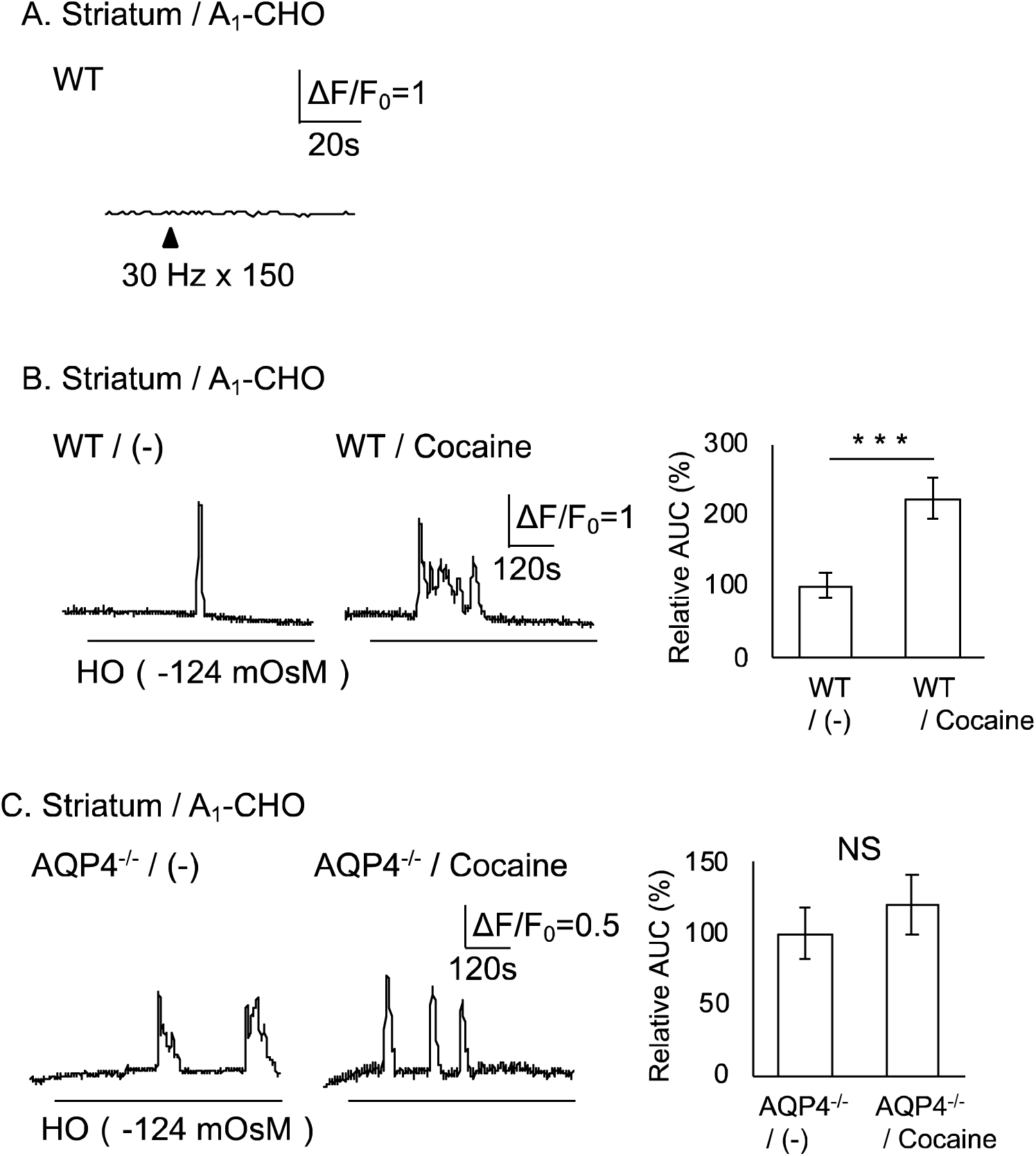
Increased adenosine release in the striatum during depressive behavior. Adenosine from striatum slices were measured by A_1_-CHO (Striatum / A_1_-CHO). A. Representative time dependent changes of [Ca^2+^]_i_ during electrical stimulation (arrowhead, 30 Hz, 150 pulses) to slices prepared from wild type mice (WT). B. Adenosine releases from slices of saline- (WT / (-), n = 6 slices) or cocaine-treated wild type mice (WT / cocaine, n = 6 slices) during hypo-osmotic treatment. Representative time dependent changes of [Ca^2+^]_i_ of A_1_-CHO (left, center). Hypo-osmotic treatment (HO; -124 mOsM) were indicated by lines. [Ca^2+^]_i_ increase during HO treatment were shown as relative area under the curves (AUC, right) normalized by control. C. Adenosine releases from slices of saline (AQP4^-/-^ / (-), n = 6 slices) or cocaine (AQP4^-/-^ / Cocaine, n = 6 slices) treated AQP4^-/-^ mice during hypo-osmotic treatment. Mean ± SE, *** p<0.001, NS, not significant.

### The suppression of dopamine release by the AQP4-dependet adenosine tone in the striatum

The influence of adenosine on the dopamine releases in the striatum was examined. An electrical stimulation (40 pulses, 30 Hz) to striatal slices induced dopamine releases, which were measured as the Ca^2+^ elevations in the D_2_-CHO. 10 µM adenosine significantly reduced the dopamine release (WT / Ado) to 61.92 ± 7.44% of control (WT / (-)) (Fig. 3A), while a non-selective adenosine receptor antagonist, 1 mM caffeine 146.67±23.50% significantly increased dopamine release (WT / Caffeine) to control (WT / (-)) (Fig. 3B). Furthermore, an A_1_ antagonist, 10 µM DPCPX 156.32±17.39% significantly increased dopamine release (WT / DPCPX) to control (WT / (-)) (Fig. 3C), while an A_2A_ receptor antagonist, 50 nM SCH58261 did not change the dopamine release (WT / SCH) from control (WT / (-)) (Fig. 3D). These results indicate that exogenous adenosine, as well as endogenous adenosine tone suppress dopamine releases via A_1_ receptor in the striatum. The involvement of AQP4 in the adenosine tone suppressing the dopamine releases in the striatum was examined using AQP4^-/-^ mice. The dopamine releases in the striatal slices prepared from AQP4^-/-^ mice (AQP4^-/-^) was 181.38±19.54% significantly larger than control (WT) (Fig. 3E). In addition, DPCPX significantly decreased the dopamine release of AQP4^-/-^ mice (AQP4^-/-^ / DPCPX) to 54.70±8.49% of control (AQP4^-/-^ / (-) (Fig. 3F). Thus DPCPX increased dopamine release in wild type mice, but did not in AQP4^-/-^ mice. These results indicates that the striatal dopamine release is suppressed by the AQP4-dependent adenosine tone and its A_1_ receptor activation.

**Fig. 3.**
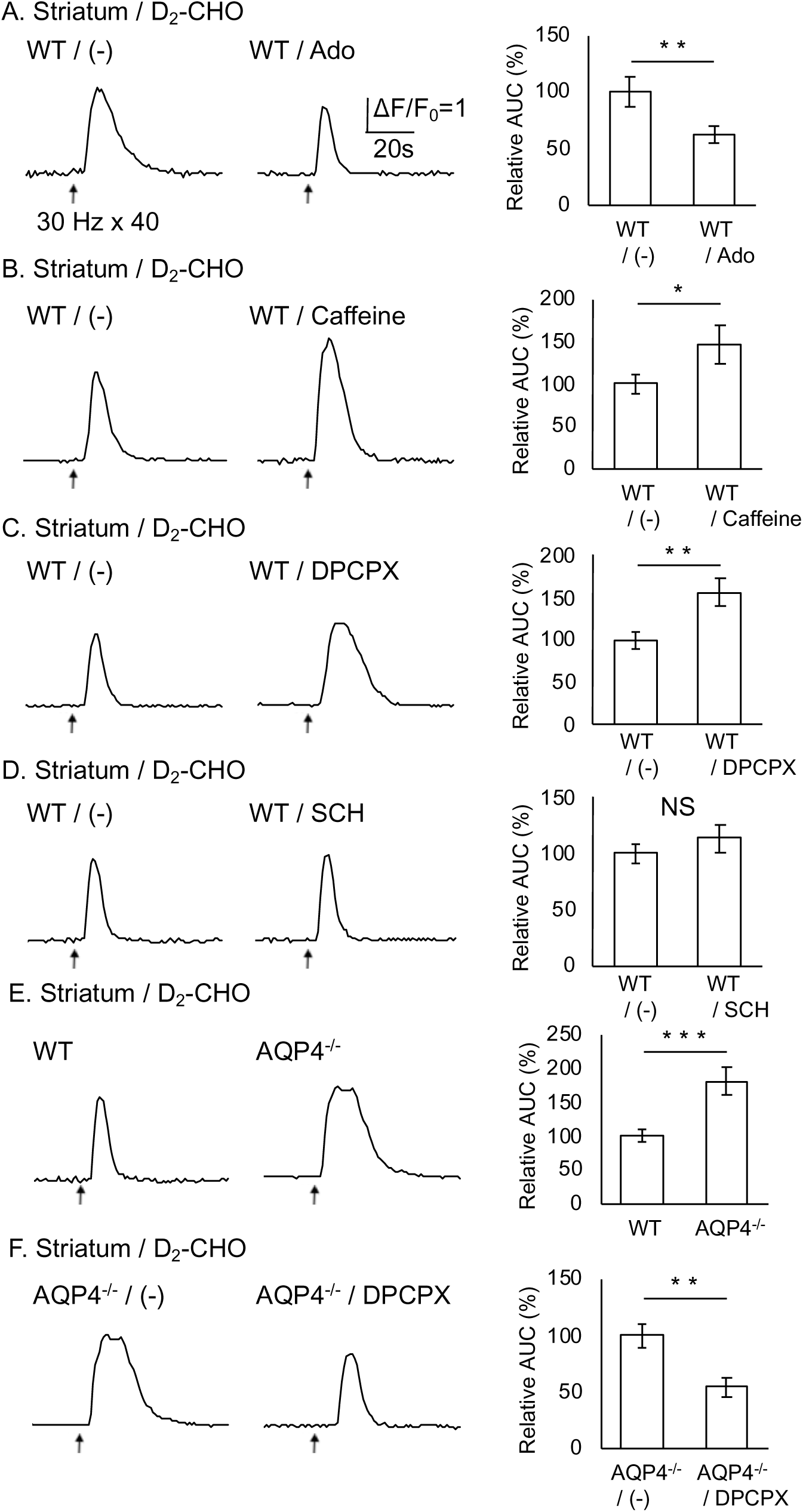
Suppression of evoked-dopamine release in the striatum by AQP4 dependent increase of adenosine. Evoked-dopamine releases in striatal slices were measured by D_2_-CHO (Striatum / D_2_-CHO). A. The effect of adenosine. Slices from wild type mice (WT) were electrically stimulated (30 Hz, 40 pulses) in the absence (WT / (-), n = 4 slices) or presence of 10 µM adenosine (WT / Ado, n = 5 slices). Representative time dependent changes of [Ca^2+^]_i_ of D_2_-CHO (left, center) during electrical stimulation (arrow). [Ca^2+^]_i_ increases of D_2_-CHO for one minute after stimulation were shown as relative AUC (right) normalized by control. B. The effect of caffein. Evoked-dopamine release in the absence (WT / (-), n = 6 slices) or presence of 1 mM caffein (WT / Caffeine, n = 6 slices). C. The effect of DPCPX. Evoked-dopamine release in the absence (WT / (-), n = 6 slices) or presence of 10 µM DPCPX (WT / DPCPX, n = 4 slices). D. The effect of an A_2A_ selective antagonist SCH58261. Evoked-dopamine release in the absence (WT / (-), n = 6 slices) or presence of 10 µM DPCPX (WT / DPCPX, n = 4 slices). E. The involvement of AQP4. Evoked-dopamine releases in wild type (WT, n = 6 slices) and AQP4 knock out mice (AQP4^-/-^, n = 7 slices). F. The effect of DPCPX on AQP^-/-^ mice. Evoked-dopamine releases in the absence (AQP4^-/-^ / (-), n = 7 slices) or presence of DPCPX (AQP4^-/-^ / DPCPX, n = 5 slices). Mean ± SE. * p<0.05, ** p<0.01, *** p<0.001, NS, not significant, t-test.

### DPCPX increased dopamine release in wild type mice, while decreased in AQP4^-/-^ mice

The further suppression of dopamine release of AQP4^-/-^ mice in the presence of DPCPX propose a facilitation of dopamine release by an A_1_ receptor subpopulation, which is activated by adenosine release through other pathway rather than the AQP4 dependent pathway as will be discussed later in mPFC results.

### Reduced dopamine release during depressive behavior

Dopamine releases in the striatum during depressive behavior was examined. The dopamine release of cocaine-treated mice (WT / Cocaine) was significantly smaller to 68.35±8.53% of control (WT / (-)) (Fig. 4A). DPCPX (WT / Cocaine / DPCPX) 137.99±18.11% significantly increased the dopamine release of cocaine-treated WT mice (WT / Cocaine / (-)) as in the case of saline-treated WT (Fig. 4B). These results suggests that the suppression of dopamine release by endogenous adenosine via A_1_ receptor persists during depressive behavior. In contrast, the dopamine release of cocaine-treated AQP4^-/-^ mice (AQP4^-/-^ / Cocaine) was not significantly different from control (AQP4^-/-^ / (-)) suggesting the ablation of reduced dopamine release by cocaine in AQP4^-/-^ mice (Fig. 4C). These results indicate the lack of depressive behavior and associated reduction of dopamine release in AQP4^-/-^ mice. In addition, DPCX did not change the dopamine release of cocaine-treated AQP4^-/-^ mice (AQP4^-/-^ / Cocaine / DPCPX and AQP4^-/-^ / Cocaine / (-)) (Fig. 4D), indicating the loss of the A_1_ receptor-mediated suppression of dopamine release in cocaine treated AQP4^-/-^ mice. From these results, cocaine is supposed to induce depressive behavior by the increase of AQP4-dependent adenosine increase and subsequent suppression of dopamine release by A_1_ receptors activation. Furthermore, A_1_ receptor antagonists likely suppressed depressive behavior by restoring adenosine tone in the striatum.

**Fig. 4.**
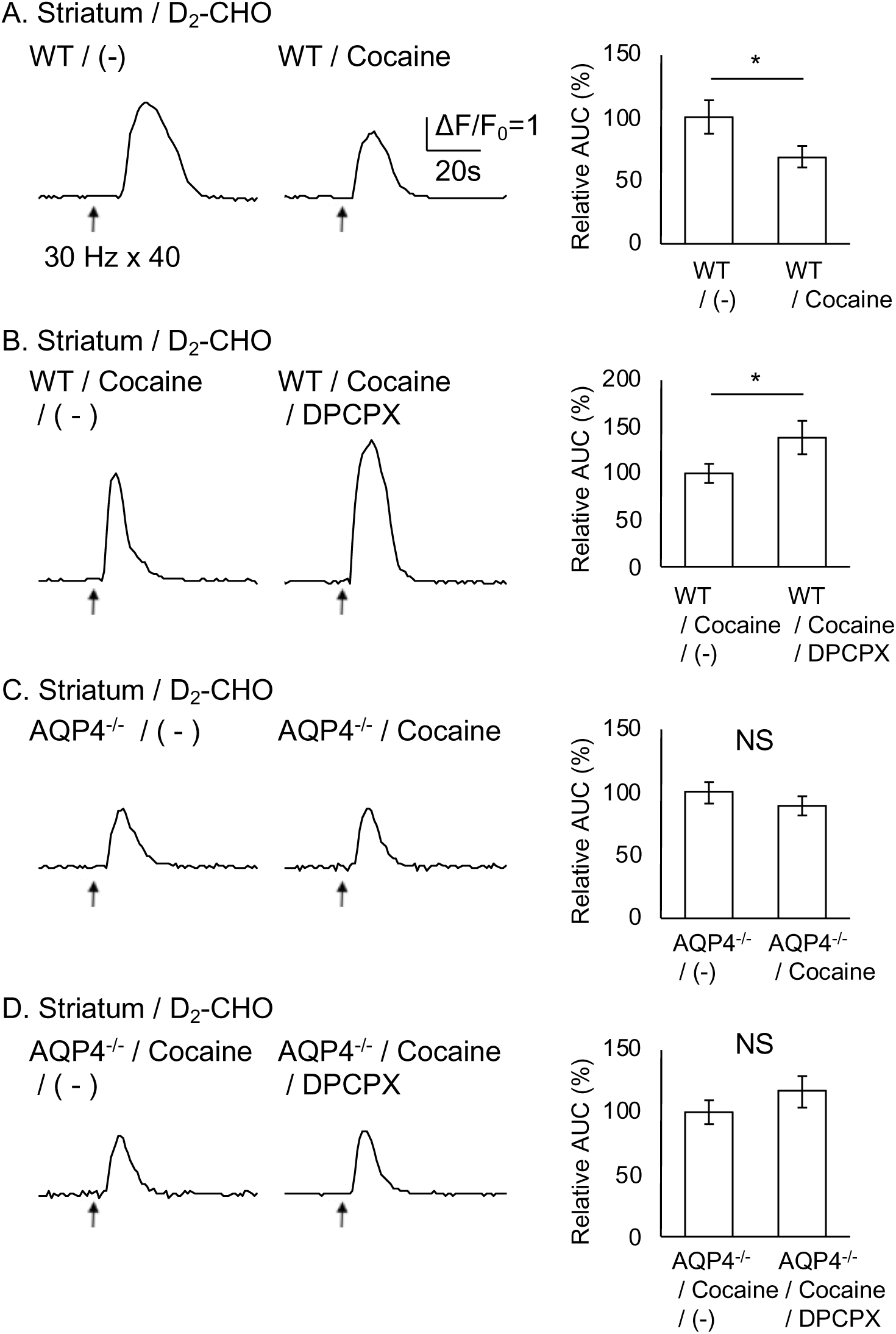
Reduction of evoked-dopamine release in the striatum of mice during depressive behavior and its reversal in AQP4 knock out mice. Dopamine releases in striatal slices were measured by D_2_-CHO (Striatum / D_2_-CHO). A. The effect of cocaine. Slices from wild type mice (WT) with (WT / cocaine, n = 7 slices) or without cocaine treatment (WT / (-), n = 10 slices) were electrically stimulated (30 Hz, 40 pulses). B. The effect of DPCPX in cocaine treated mice. Evoked-dopamine release in the absence (WT / Cocaine / (-), n = 6 slices) or presence of 1 mM caffein (WT / Cocaine / DPCPX, n = 5 slices). C. The effect of cocaine on AQP4^-/-^ mice. Evoked-dopamine releases in AQP4^-/-^ mice with (AQP4^-/-^ / cocaine, n = 9 slices) or without cocaine treatment (AQP4^-/-^ / (-), n = 9 slices). D. The effect of DPCPX in cocaine-treated AQP4^-/-^ mice. Evoked-dopamine release in cocaine-treated AQP4^-/-^ mice in the absence (AQP4^-/-^ / Cocaine /(-), n = 7 slices) or presence of cocaine (AQP4^-/-^ / Cocaine / DPCPX, n = 6 slices). Mean ± SE. * p<0.05, NS, not significant, t-test.

### Adenosine releases in mPFC

Since the altered neural activities in the mPFC is known to cause depressive behavior (Furuyashiki, 2012), the AQP4-dependent adenosine release in the mPFC was analyzed by using A_1_-CHO. Both 150 pulses electrical stimulation at 30 Hz and -82.7 mOsM hypo-osmotic treatment induced adenosine releases in mPFC slices. The AUC after electrical stimulation did not show significant difference between AQP4^-/-^ mice (AQP4^-/-^) and control (WT) (Fig. 5A). In contrast, hypo-osmotically-induced adenosine release of AQP4^-/-^ mice (AQP4^-/-^) was 64.14±9.24% significantly reduced to control (WT) (Fig. 5B). These results indicate that adenosine release in the mPFC is in line with our previous study using hippocampal slices, where both electrical stimulation and hypo-osmotic treatment induce adenosine release and the hypo-osmotic release depends on AQP4 (Yamashiro et al., 2017). In order to determine the involvement of mPFC adenosine in depressive behavior, adenosine releases in cocaine-treated mice were examined. As results, both electrically and hypo-osmotically-induced adenosine release of cocaine-treated mice (WT / Cocaine) did not significant difference from those of control (WT / (-)) (Fig. 5C,D). Thus, it was suggested that these adenosine releases in the PFC were not involved in depressive behavior.

**Fig. 5.**
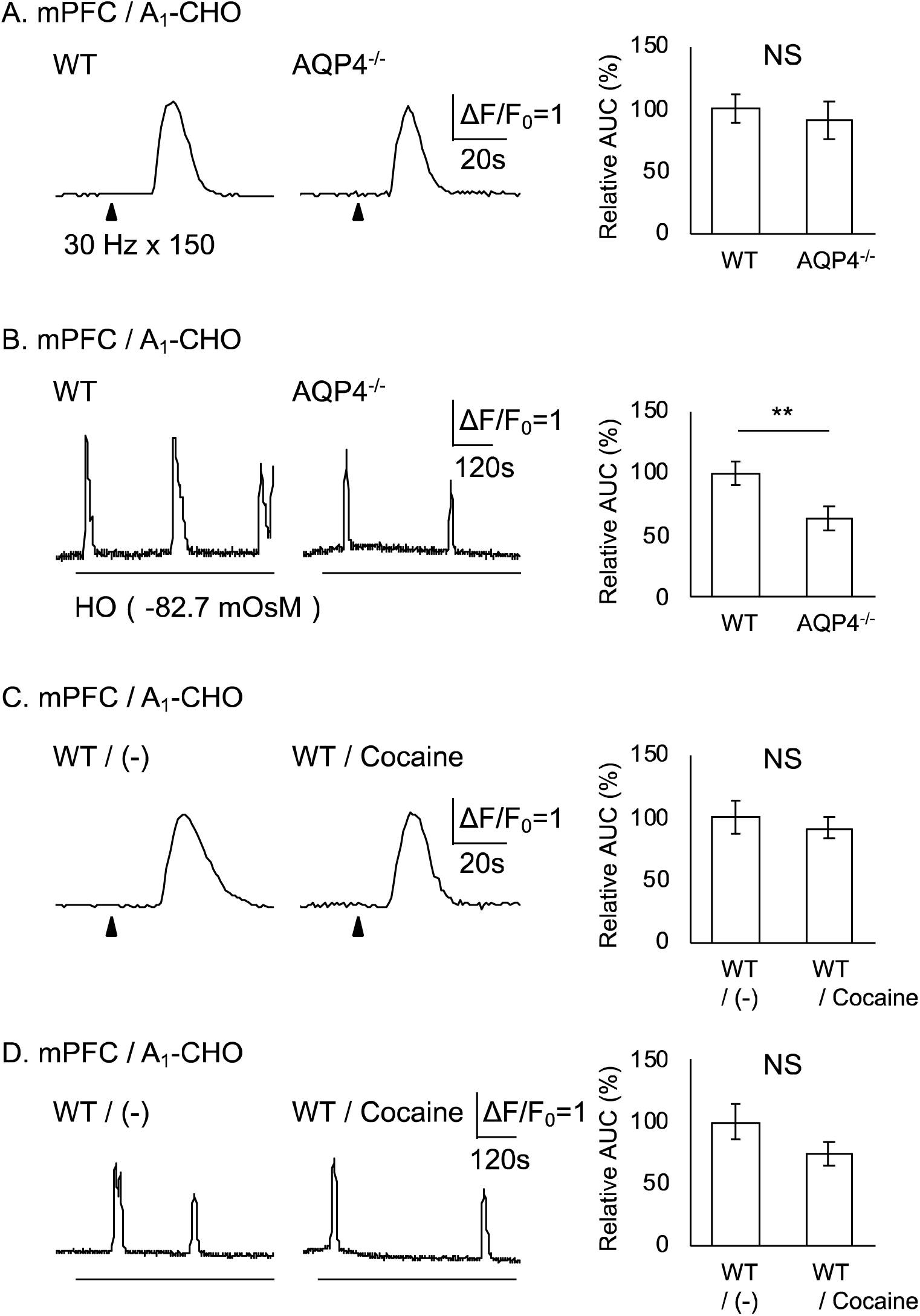
Adenosine release in mPFC is not associated with depressive behavior. Adenosine from medial prefrontal cortex (mPFC) slices were measured by A_1_-CHO (mPFC / A_1_-CHO). A. Evoked-adenosine releases from wild type (WT, n = 6 slices) and AQP4^-/-^ mice (AQP4^-/-^, n = 5 slices). Representative time dependent changes of [Ca^2+^]_I_ of A_1_-CHO (left, center) during electrical stimulation (150 pulses at 30 Hz, arrowhead). [Ca^2+^]_i_ increases of A_1_-CHO for one minute after stimulation were shown as relative AUC (right) normalized by control. B. Slices from wild type mice (WT, n = 6 slices) and AQP4^-/-^ mice (AQP4^-/-^, n = 6 slices) were hypo-osmotically stimulated (HO; -82.6 mOsM) as indicated by lines. C. Evoked-adenosine releases in the absence (WT / (-), n = 7 slices) or presence of cocaine (WT / Cocaine, n = 6 slices. D. Hypo-osmotically-induced adenosine release in mice with (WT / cocaine, n = 6 slices) or without cocaine treatment (WT / (-), n = 6 slices). Mean ± SE, ** p<0.01, NS, not significant.

### The involvement of adenosine in dopamine release in the mPFC

Since the decreased dopamine neurotransmission in the mPFC during depressive behavior is proposed (Furuyashiki, 2012), the involvement of adenosine in evoked-dopamine release in the mPFC was examined by using D_2_-CHO. As in the striatum, the dopamine releases were induced by 40 pulses electrical stimulation at 30 Hz (arrow) in the mPFC. Exogenous adenosine 67.12±8.13% significantly reduced dopamine release in the mPFC (WT / Ado) to control (WT / (-)) (Fig. 6A). Meanwhile, caffeine 66.96±5.82% significantly reduced in dopamine release in the mPFC (WT / Caffeine) to control (WT / (-)) (Fig. 6B). In addition, DPCPX 75.25±5.45% significantly reduced dopamine release (WT / DPCPX) to control (WT / (-)) (Fig. 6C), while SCH58261 did not change dopamine release (WT / SCH) to control (WT / (-)) (Fig. 6D). These results demonstrate the suppression of dopamine release in the PFC both by exogenous adenosine and A_1_ receptor that the A_1_ receptor activation by endogenous adenosine enhance dopamine release as in the opposite case in the striatum.

**Fig. 6.**
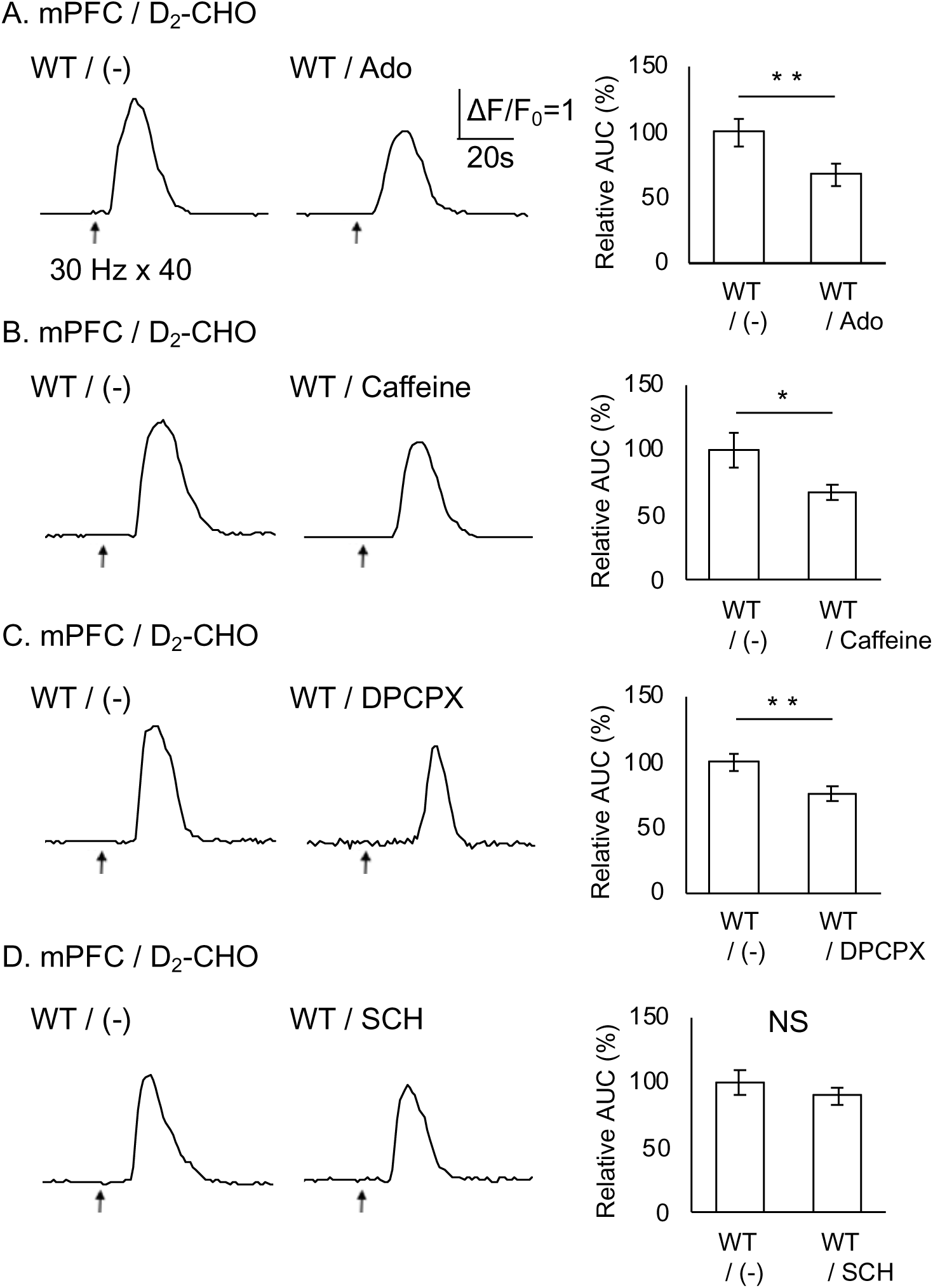
Suppression of evoked-dopamine release in the mPFC by adenosine. Dopamine from mPFC slices were measured by D_2_-CHO (mPFC / D_2_-CHO). A. The effect of adenosine. Slices from wild type mice (WT) were electrically stimulated (30 Hz, 40 pulses) in the absence (WT / (-), n = 8 slices) or presence of adenosine (WT / Ado, n = 9 slices). B. The effect of caffein. Evoked-dopamine releases in the absence (WT / (-), n = 8 slices) or presence of caffein (WT / Caffeine, n = 8 slices). C. The effect of DPCPX. Evoked-dopamine releases in the absence (WT / (-), n = 8 slices) or presence of DPCPX (WT / DPCPX, n = 9 slices). D. The effect of SCH58261. Evoked-dopamine releases in the absence (WT / (-), n = 10 slices) or presence of SCH58261 (WT / SCH, n = 10 slices). Mean ± SE. * p<0.05, ** p<0.01, NS, not significant, t-test.

### Downregulation of A_1_ receptor in the mPFC during depressive behavior

Dopamine releases in the mPFC during depressive behavior were examined by D_2_-CHO. The electrically induced dopamine releases in the mPFC slices from cocaine treated mice (WT / Cocaine) was not significantly different from control (WT / (-)) (Fig. 7A). This suggests the dopamine release in mPFC was not affected by the cocaine. However, DPCPX, which increased the dopamine releases in the mPFC of naïve mice, did not change the dopamine release in cocaine treated mice (WT / Cocaine / DPCPX and WT / Cocaine / (-)), suggesting the loss of the DPCPX induced increases of dopamine releases by cocaine (Fig. 7B). Since cocaine did not change adenosine release in the mPFC, the failure of DPCPX to change dopamine release in cocaine-treated mPFC is supposed to reflect downregulation of A_1_ receptors. This point was examined in AQP4 AQP4^-/-^ mice. Cocaine treatment did not change dopamine release in the mPFC of AQP4^-/-^ mice ( (AQP4^-/-^ / (-) and AQP4^-/-^ / Cocaine) (Fig. 7C). Meanwhile, DPCPX 61.26±5.27% significantly reduced dopamine release of cocaine-treated AQP4^-/-^ mice (AQP4^-/-^ / Cocaine / DPCPX) to control (AQP4^-/-^ / Cocaine / (-)) (Fig. 7D). Hence, both the depressive behavior and the down-regulation of A_1_ receptors were lost in AQP4^-/-^ mice. All together, these results suggest that cocaine induces depressive behavior by down-regulating A_1_ receptor in the mPFC through an AQP4-dependent pathway.

**Fig. 7.**
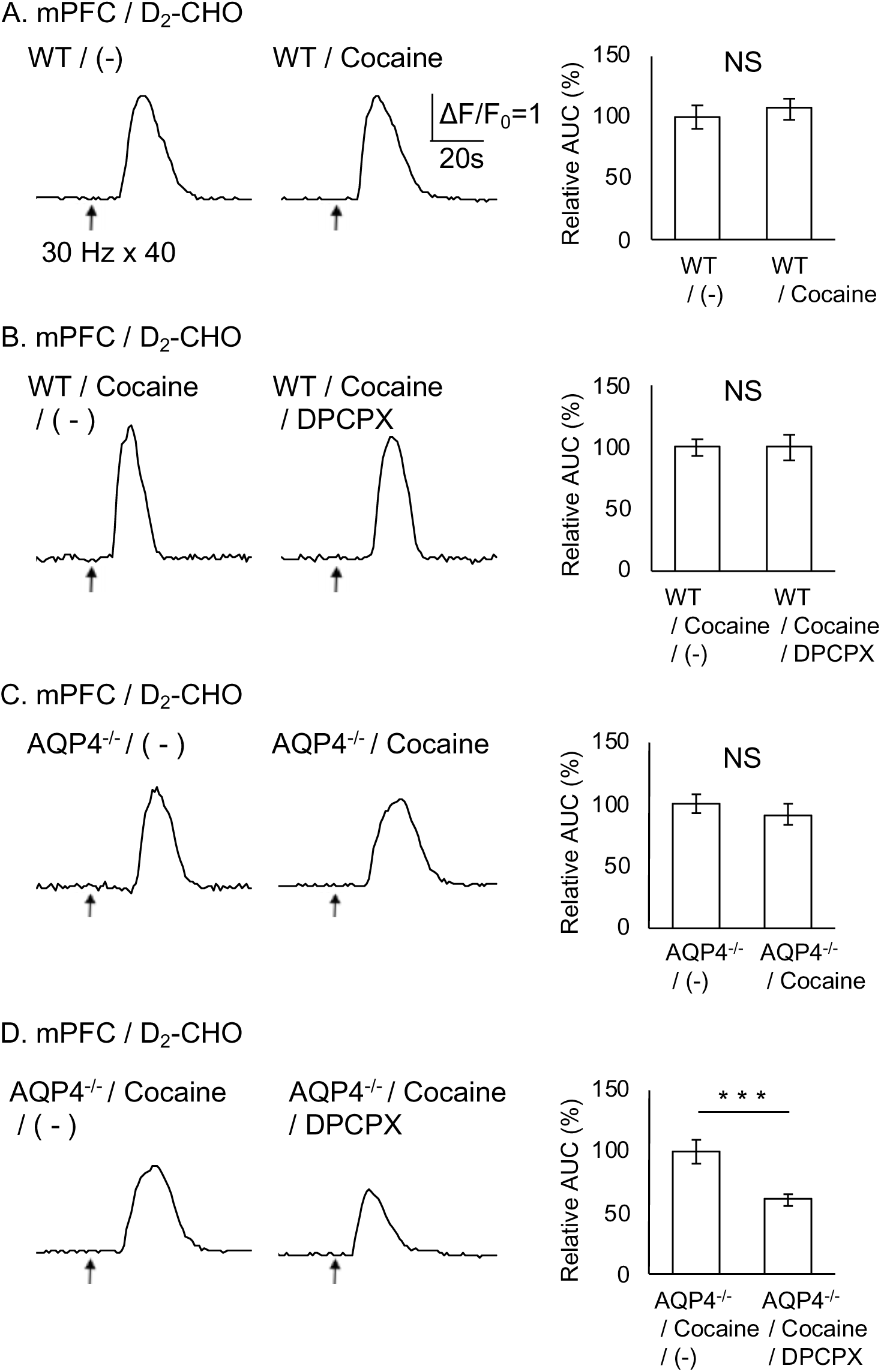
Downregulation of A_1_ receptor in the mPFC of cocaine-treated mice. Dopamine from mPFC slices were measured by D_2_-CHO (mPFC / D_2_-CHO). A. The effect of cocaine. Slices from wild type mice (WT) with (WT / cocaine, n = 12 slices) or without cocaine treatment (WT / (-), n = 9 slices) were electrically stimulated (30 Hz, 40 pulses). B. The effect of DPCPX in cocaine treated mice. Evoked-dopamine releases in the absence (WT / Cocaine / (-), n = 8 slices) or presence of DPCPX (WT / Cocaine / DPCPX, n = 8 slices). C. The effect of cocaine in AQP4^-/-^ mice. Evoked-dopamine releases in absence (AQP4^-/-^ / (-), n = 9 slices) or presence of cocaine (AQP4^-/-^ / cocaine, n = 8 slices). D. The effect of DPCPX in cocaine treated AQP4^-/-^ mice. Evoked-dopamine releases in the absence (AQP4^-/-^ / Cocaine /(-), n = 8 slices) or presence of DPCPX (AQP4^-/-^ / Cocaine / DPCPX, n = 7 slices). Mean ± SE. * p<0.05, NS, not significant, t-test.

### AQP4-dependent adenosine tone facilitates dopamine release in the mPFC

The involvement of AQP4-dependent adenosine tone in evoked-dopamine release in the mPFC was examined by measuring the evoked-dopamine release of AQP4^-/-^ mice using D_2_-CHO. The dopamine release in the mPFC of AQP4^-/-^ mice (AQP4^-/-^) was not significantly different from wild type mice (WT) (Fig. 8A). The dopamine release in the mPFC of AQP4^-/-^ mice in the presence of DPCPX, which decrease the dopamine release in mPFC of WT mice was also not different from control (AQP4^-/-^ / (-)) (Fig. 8B). These results indicate the lack of A_1_ receptor-mediated reduction of dopamine release in the mPFC of AQP4^-/-^ mice. Since hypo-osmotically-induced adenosine increase was smaller in the mPFC of AQP^-/-^ mice, the loss of A_1_ receptor-mediated reduction of dopamine release is likely due to the loss of A_1_ receptor rather than the change of AQP4-dependent adenosine tone.

**Fig. 8.**
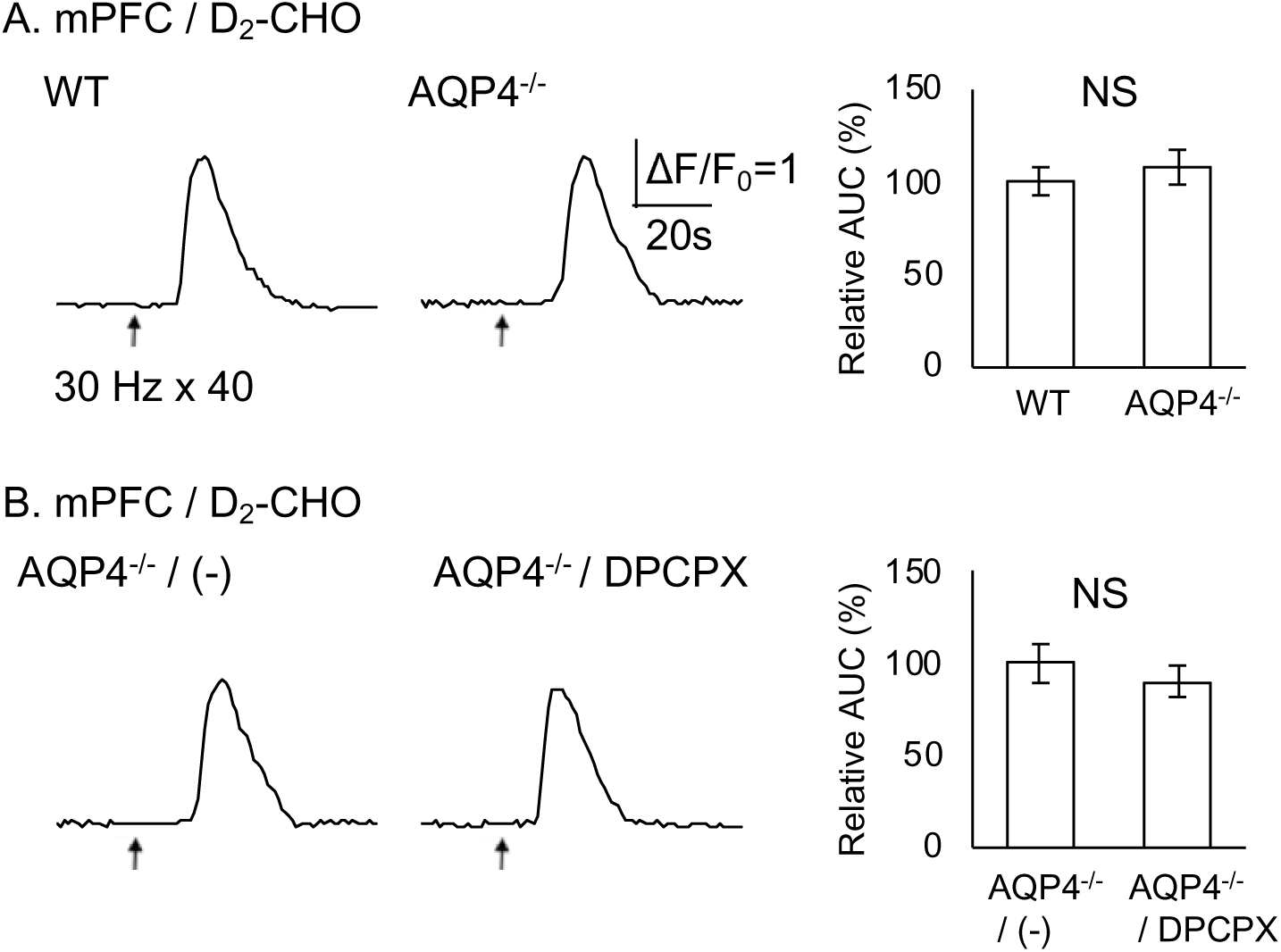
Facilitation of evoked-dopamine release in the mPFC by AQP4-dependent increase of adenosine. Evoked-dopamine release from mPFC slices were measured by D_2_-CHO (mPFC / D_2_-CHO). A. The involvement of AQP4. Slices from wild type mice (WT, n = 7 slices) and AQP4 knock out mice (AQP4^-/-^, n = 7 slices) were electrically stimulated (30 Hz, 40 pulses). B. The effect of DPCPX on AQP4^-/-^ mice. Evoked-dopamine releases in the absence (AQP4^-/-^ / (-), n = 7 slices) or presence of DPCPX (AQP4^-/-^ / DPCPX, n = 8 slices). NS, not significant, t-test.

### Two adenosine-dependent pathways regulating dopamine release

The present discrepancy that both exogenous adenosine and DPCPX reduce dopamine release in mPFC was further examined. A_1_ receptor is commonly involved in pre-synaptic inhibition, but the inhibition of A_1_ receptor suppressed dopamine release in the mPFC. Pre-synaptic inhibition of dopamine release is mediated not only by A_1_ receptor, but also other receptors, such as metabotropic glutamate receptor (*) or GABA_A_ receptor (Yonezawa et al., 1998). Thus, one possibility is that a subpopulation of A_1_ receptor inhibits glutamate or GABA release, which are simultaneously activated with dopamine release by electrical stimulation and pre-synaptically inhibit dopamine release.

To test this possibility, the effects of DPCPX in the presence of mGluR or GABA_A_ receptors antagonist were examined. The effect of DPCPX was examined in the presence of GABA_A_ receptor antagonist, 100 µM picrotoxin. In the presence of picrotoxin and DPCPX, the dopamine release (WT / PTX / DPCPX) was not significantly different from the control (WT / PTX / (-)) (Fig. 9A). Therefore, GABA_A_ receptor antagonist occlude the effect of DPCPX. Since A_1_ receptors are mostly resided in excitatory synapses rather than inhibitory ones (Chen et al., 2013), DPCPX supposed to affect on glutamatergic synapses. Since Group II metabotropic glutamate receptors (Group II mGluRs) regulate dopamine release in the mPFC and nucleus accumbens (Cartmell et al., 2000; Gupta & Young, 2018), the possibility that DPCPX affects glutamatergic terminal involving Group II mGluR was examined.

**Fig. 9.**
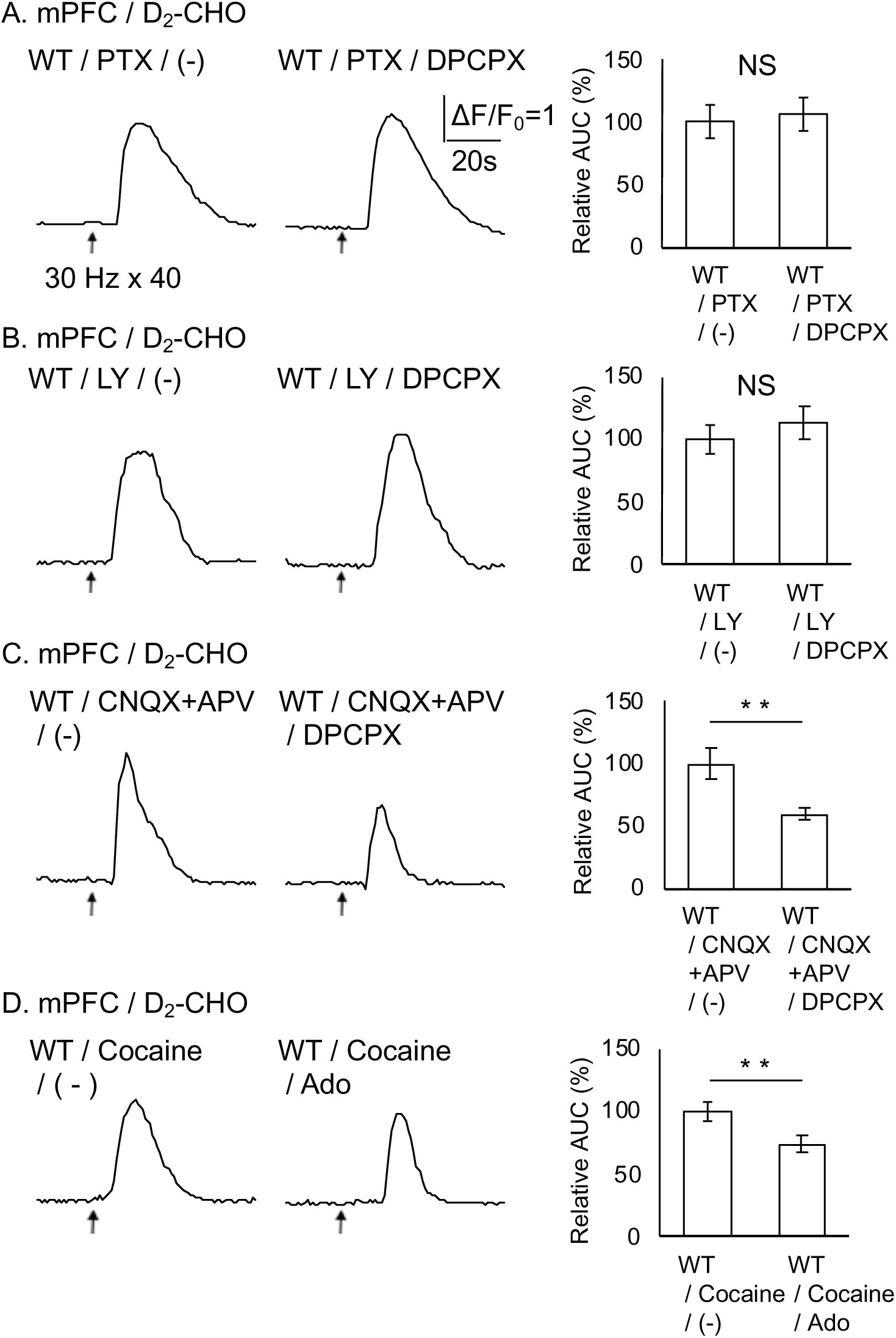
Dual regulation of dopamine release in the mPFC by adenosine tone. Evoked-dopamine releases from mPFC slices were measured by D_2_-CHO (mPFC / D_2_-CHO). A. The effect of DPCPX in the presence of a GABA receptor antagonist picrotoxin, PTX. Slices from wild type mice (WT) were electrically stimulated (30 Hz, 40 pulses) in the presence of 100 µM PTX and DPCPX (WT / PTX / DPCPX, n = 6) or PTX alone (WT / PTX / (-), n = 6 slices). B. The effect of Group II mGluR antagonist LY341495 (LY). Evoked-dopamine release int the presence of 500 nM LY and DPCPX (WT / LY / DPCPX, n = 8 slices) or LY alone (WT / LY / (-), n = 8 slices). C. The effect of ion channel type glutamate receptor antagonists, CNQX and APV. Evoked-dopamine releases in the presence of 10 µM CNQX, 50 μM APV and DPCPX (WT / CNQX+APV / DPCPX, n = 9 slices) or CNQX+APV alone (WT / CNQX+APV / (-), n = 10 slices). D. The effect of adenosine on cocaine-treated mice. Evoked-dopamine releases in slices from cocaine-treated wild type mice in the absence (WT / Cocaine / (-), n = 9 slices) of presence of adenosine (WT / Cocaine / Ado, n = 9 slices). NS, not significant, t-test

The group II mGluR antagonist LY341495 was used to test this hypothesis and the dopamine release in the presence of 500 nM LY341495 and DPCPX and control (WT / LY / DPCPX and WT / LY / (-)) did not show significant difference (Fig. 9B). Hence, group II mGluR antagonist also occluded the effect of DPCPX on the dopamine release. If the DPCPX-sensitive glutamate release acts on ionotropic glutamate receptor and influences on dopaminergic neuron via neural circuit, AMPA receptor antagonist CNQX and NMDA receptor antagonist APV may affect. However, the dopamine release in the presence of DPCPX, together with 10 µM CNQX and 50 µM APV (WT / CNQX+APV / DPCPX) was 59.82 ± 4.12% significantly reduced to control (WT / CNQX+APV / (-)) (Fig. 9C), excluding the involvement of ionotropic glutamate receptors. These results indicate that the DPCPX affects the glutamate and GABA releases, which act on group II mGLuR or GABA_A_ receptor locating at the presynaptic terminal releasing dopamine, respectively. Finally, the A_1_ receptor down-regulation by cocaine treatment at dopamine release terminal, which is sensitive to exogenous adenosine in the mPFC. The dopamine release of cocaine treated mice in the presence of exogenous adenosine (WT / Cocaine / Ado) 73.79±6.82% significantly reduced to control (WT / Cocaine / (-)) (Fig. 9D). This result reflects that cocaine occluded the effect of DPCPX while did not influence on effect of exogenous adenosine. Overall, the dopamine release in the mPFC were regulated by two adenosine-dependent pathways, one pathways involving glutamate and GABA was down-regulated bycocaine treatment, while another direct adenosine inhibition of dopamine releasing terminal was not affected.

## Discussion

Pathological changes in adenosine and dopamine release associated with depressive-like behavior during cocaine withdrawal has been investigated. Depressive-like behavior accessed by the FST suggested that AQP4-mediated increase in adenosine causes depressive-like behavior via A_1_ receptors, because genetic disruption of AQP4 and pharmacological inhibition of A_1_ receptor suppressed the depressive-like behavior. Adenosine release in the striatum was not detected upon electrical stimulation while was detected by a hypoosmotic treatment, but it required greater osmotic change than in other brain regions, and was not affected by disruption of the AQP4 gene. This indicates the difference of extracellular adenosine dynamics between in the striatum, where A_2A_ receptors are highly expressed (*), and in the cortex and hippocampus. In contrast, adenosine release in the mPFC, adenosine was induced by electrical stimulation and hypoosmotic treatment, and adenosine release upon hypoosmotic treatment was AQP4 dependent, similar to our previous study on the hippocampus (Yamashiro et al., 2017). In contrast, dopamine release upon electrical stimulation was detected in both the striatum and mPFC. In the striatum, dopamine release was inhibited by adenosine, whereas it was enhanced by A_1_ receptor antagonist, suggesting tonic inhibition of dopamine release by adenosine. Furthermore, cocaine increased adenosine release while decreased dopamine release, indicating that cocaine promoted the tonic inhibition of dopamine release by adenosine in the striatum. In contrast, both adenosine and an A_1_ receptor antagonist suppressed dopamine release in the mPFC, and cocaine treatment abolished only the effect of A_1_ receptor antagonist in a AQP4-dependent manner. Thus, cocaine likely down-regulated A_1_ receptors by an AQP4-dependent mechanism in the mPFC, resulting in loss of endogenous adenosine modulation of dopamine release. Since the effects of A_1_ receptor antagonist were occluded by GABA_A_ or gruop II mGluR antagonists, A_1_ receptor antagonist likely acted on presynaptic terminal releasing GABA or glutamate, which are known to suppresses dopamine release via GABAA receptor or group II mGluR (*). These results demonstrate modulations of dopamine neurotransmission by AQP4 and adenosine in both the striatum and mPFC, and their involvement in depressive-like behavior. However, depression-like behavior during cocaine withdrawal more likely attributed to the change in the striatum, where A_1_ antagonist restored dopamine release as in the case of depression-like behavior, in contrast to the suppression of dopamine release by both exogenous adenosine and A_1_ antagonist in the mPFC.

The mitigation of depression-like behavior during cocaine withdrawal in AQP4_-/-_ mice and the suppression of dopamine release by AQP4-dependent increase in adenosine in the striatum are supported by a preceding report of increased dopamine concentrations in the striatum as measured by microdialysis and associated cognitive and psychiatric changes in AQP4_-/-_ mice (Szu and Binder, 2016). The antidepressant effects of caffeine and DPCPX against depression-like behavior during cocaine withdrawal is consistent with previously reported antidepressant effects of these compounds in a forced swimming test using normal mice (Szopa et al., 2016) (Szopa et al., 2018). The present study is the first demonstration of crosstalk between AQP4-dependent increase in adenosine and pathological change of dopamine neurotransmission in depression by measuring adenosine and dopamine releases and analyzing their changes during cocaine withdrawal.

The failure of detecting adenosine release upon electrical stimulation in striatal slices is consistent with a report of low detection success rates in the measurement of similarly-evoked adenosine release in slices by fast scan cyclic voltammetry (Pajski and Venton, 2013). Meanwhile, evoked-adenosine releases in the striatum *in vivo* were reported by using microdialysis (Cechova and Venton, 2008; Cechova et al., 2010). One possible explanation for the failure in slice is depletion of neuronal adenosine source due to high oxygen and glucose condition for slices. Another explanation is that AQP4-dependent increase in adenosine is induced by electrical stimulation *in vivo*, but not in slices lacking blood flow, which is likely essential for water exchange via AQP4 locating at the interface between astrocytes and blood vessels. We suppose that neuronal activities induce water influx from blood flow to astrocyte via AQP4, because they increase glucose uptake or glycogen degradation, both of which increase osmotic pressure in astrocytes. Adenosine release in striatal slices was induced by hypoosmotic treatment at -120 mOsM, but not at -82.7 mOsM, which was effective in our previous study using hippocampal slices (*) and the present study using mPFC slices. In addition, AQP4-/- mice showed reduction of hypoosmotically-induced adenosine release in cocaine treated mice, but not in normal ones. Thus, it is supposed that astrocytes are capable of releasing ATP even without AQP4 following intense change of osmolarity, and the availability of AQP4 in the striatum is upregulated by cocaine treatment. We have been investigating changes of astrocyte structure and AQP4 expression in the striatum during cocaine withdrawal, however no significant result is obtained.

Dopamine release in the striatum was suppressed by exogenous adenosine, while increased by DPCPX and in AQP4^-/-^ mice. These results indicates the tonic suppression of dopamine release in the striatum by adenosine released through AQP4-dependent pathway. This point is consistent with previous reports showing presynaptic inhibition of dopamine release by A_1_ receptor (Yabuuchi et al., 2006), but not by A_2A_ receptor (Ross and Venton, 2015). In the present study, DCPCX significantly reduced evoked-dopamine release in AQP4-/- striatum. This result can be interpreted as the suppressive effect of DPCPX in the mPFC, namely, adenosine release by an AQP4-indeoendent pathway in the striatum presynaptically suppresses other neuromodulator, such as GABA and glutamate, which negatively regulate dopamine release.

